# Negative association between nodular stromal fibrosis and large lipid droplets in human choroid plexus epithelial cells

**DOI:** 10.64898/2026.11.12.766290

**Authors:** Michael J. Neel, N. Victoria Espericueta, Aarij Gora, Jessica M. Yeung, Ashley A. Nguyen, Juslyn Z. Chan, William Huang, Sam Ghahremani, Eunice Shin, Jonathan Zebroski, Jackson Sousa, Nathan Schacherl, Kean Ehsani, Christina M. Magana-Ramirez, Daniel Gillen, Peter Chang, Brett A. Johnson, Edwin S. Monuki

**Affiliations:** Department of Pathology & Laboratory Medicine, University of California, Irvine; Department of Statistics, University of California, Irvine, California, USA; Institute for Memory Impairments and Neurological Disorders, University of California, Irvine, California, USA; Department of Epidemiology and Biostatistics, University of California, Irvine, California, USA; Department of Population Health and Disease Prevention, University of California, Irvine, California, USA; Alzheimer’s Disease Research Center, University of California Irvine, Irvine, CA, USA; Department of Radiological Sciences, University of California, Irvine, Irvine, CA, USA; Donald Bren School of Information and Computer Sciences, University of California Irvine, Irvine, CA, USA; Center for Artificial Intelligence in Diagnostic Medicine, University of California Irvine, Irvine, CA, USA; Department of Developmental & Cell Biology, University of California Irvine, Irvine, CA, USA; Sue and Bill Gross Stem Cell Research Center, University of California Irvine, Irvine, CA, USA

## Abstract

Fibrosis is an age-related change within the human choroid plexus that is likely to affect its function negatively. We systematically analyzed nodular stromal fibrosis present in postmortem tissue from a large repository encompassing the human lifespan. Following manual annotation of sampled images from 81 specimens of choroid plexus from the atrium of the lateral ventricle, a deep learning model was trained to assess stages of fibrosis (dense, hyalinized, and mineralized) occurring in papillae across whole slide images of hematoxylin and eosin-stained sections from 204 cases. The percentage of affected papillae and the relative area of fibrosis at all three stages of severity increased with age. Affected papillae were significantly clustered across the choroid plexus, with areas of high and low fibrotic density. Choroid plexus epithelial cells (CPECs) associated with fibrotic papillae were less likely to contain large lipid droplets than were non-fibrotic papillae, as assessed by another deep learning model. Across papillae within a case, the area of each stage of fibrosis was negatively correlated with the prevalence of CPECs containing large lipid droplets. Finally, individuals displaying more nodular fibrosis had significantly fewer CPECs containing large lipid droplets. Thus, age-related nodular stromal fibrosis may negatively impact lipid storage and homeostasis for the choroid plexus and brain.

## INTRODUCTION

Fibrosis, an excessive accumulation of extracellular matrix components, affects numerous tissues and is associated with multiple diseases (Antar et al 2023, Rieder et al 2025). Inflammation following repeated injury can result in fibrotic scarring that alters tissue structure and impairs vital organ functions (Antar et al 2023, Rieder et al 2025). Fibrosis has long been known to affect the aged human choroid plexus (ChP) (Taft 1922, Dunn & Kernohan 1955, Shuangshoti & Netsky 1970, Prineas et al 2016, Xu et al 2026), the papillary tissue that provides important homeostatic support for the central nervous system by producing and conditioning the cerebrospinal fluid (CSF), creating the blood-CSF barrier, and providing a pathway for immune cell entry into the brain (Lehtinen et al 2013, Lun et al 2015, Spector et al 2015, Ghersi-Egea et al 2018).

In the human ChP, fibrosis has been associated with expansion of the stroma (Dunn & Kernohan 1955), thinning of the overlying epithelium (Dohrmann & Bucy 1970), stromal mineralization (Taft 1922, Dunn & Kernohan 1955, Shuangshoti & Netsky 1970, Dohrmann & Bucy 1970), loss of capillaries within papillary stromal cores (Taft 1922, Prineas et al 2016), and the presence of complement proteins, inflammatory cells, and IgG (Prineas et al 2016). The observed changes to ChP structure associated with fibrosis suggest pathological consequences both to the tissue and to the central nervous system it supports.

We recently have found that human ChP epithelial cells (CPECs) contain large (> 5 micron) lipid droplets that are characteristic of cells such as hepatocytes, hepatic stellate cells, enterocytes, adipocytes (all of which supply lipids to other organs), and mammary epithelial cells (which supply milk fat for offspring), and we have proposed that CPECs may store lipid for use by the central nervous system as well as for their own extensive energy demands (Espericueta et al 2026). The percentage of postmortem CPECs containing large lipid droplets was correlated with age and body mass index at death (Espericueta et al 2026). Consistent with the relationship between adiposity and postmortem large lipid droplets, human stem cell-derived CPECs accumulated large lipid droplets when fatty acids were added to the medium (Espericueta et al 2026).

In the present study, we expand our analysis of age-related changes affecting the human ChP (Espericueta et al 2026) to include a systematic analysis of nodular fibrosis using our large repository of postmortem ChP tissue extracted from the atrium of the lateral ventricle. We have employed deep-learning (DL) artificial intelligence models to classify stages of fibrosis (dense, hyalinized, and mineralized), which we express both as percentages of papilla and as percentages of papillary area affected. We characterize the accumulation of fibrosis with age, find that fibrotic papillae are spatially clustered within the ChP, and discover that there is a negative association between degrees of fibrosis and percentages of CPECs containing large lipid droplets. These results suggest that fibrosis may disrupt lipid homeostasis for the ChP and brain.

## METHODS

### Tissue

Lateral ventricle ChP (glomus) was taken from the atrium/trigone during cutting of fixed brains by either the UCI Alzheimer’s Disease Research Center (ADRC) or the UCI Medical Center autopsy service. In addition to differences in postmortem intervals (ADRC < MC), autopsy service brains were fixed in formalin, while ADRC brains were fixed in freshly prepared 4% paraformaldehyde. Specimens were embedded in paraffin, and 5-µm sections were stained using hematoxylin and eosin (H&E) by the UCI Experimental Tissue Resource. Stained sections were scanned into whole slide images at 40X using an Aperio Versa 2 scanner in the ADRC neuropathology core. Image files were converted to svs and HDF5 formats compatible with a custom software client for manual annotation and neural network development.

### Manual annotations

The software client subdivided whole slide images into tiles of 561 x 561 µm with a buffer of 164 µm on each side, and these were presented to annotators in random order, with a stereological frame superimposed. Annotators traced qualified papillae between bordering concavities along the outer boundary of their stroma (Fig.1). Annotators then separately traced any areas within the papillae that were densely fibrotic (eosinophilic areas where fibers coalesced to no longer be individually distinct, but with intermittent unstained patches), hyalinized (glassy appearance with no unstained patches), and/or mineralized (concretions that stain intensely purple by H&E) (Fig. 1).

**Fig. 1.**
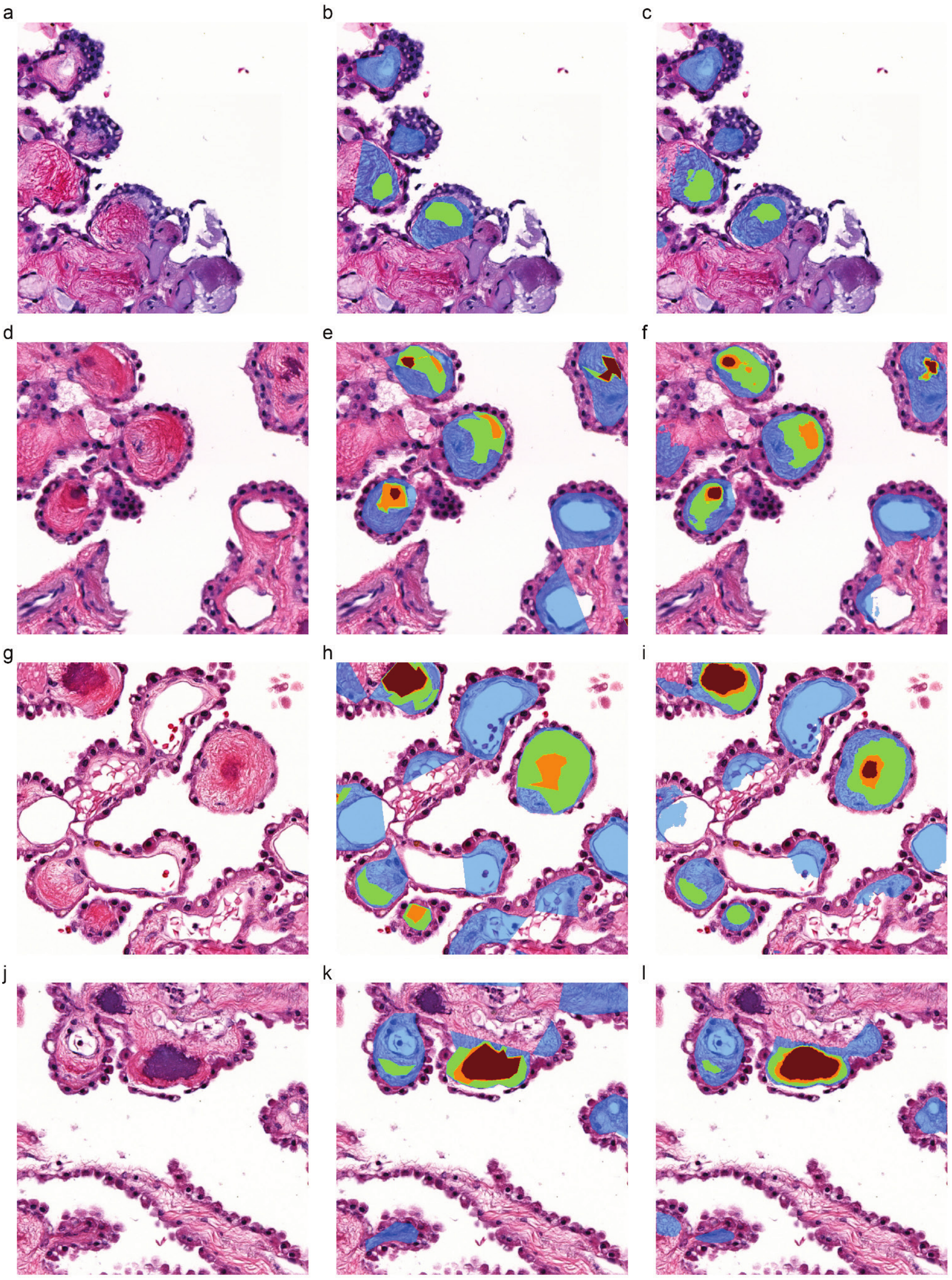
DL model improves upon manual fibrosis segmentations. H&E-stained images of fibrosis showing no annotations (**left**), manual fibrosis segmentations (**center**), and DL-model generated segmentations (**right**). Segmentations produced by the DL-model appear smoother and more accurate in contouring papillae and fibrotic regions. Transparent blue regions represent papillary areas. Green, orange, and dark red segmentations are dense, hyalinized, and mineralized fibrotic areas, respectively.

The software client calculated and plotted the percent affected area and coefficient of error (COE) over the previous ten tiles for each category of fibrosis. Once COE was <5% for 10 continuous tiles for all categories exceeding 2% of papilla area, annotators were prompted to export results. Given these criteria, at least 20 tiles were annotated per case, and on average, the <5% COE threshold was met after annotating 850 papilla. Any tiles lacking papilla were ignored in calculating the completion criteria. This sampling method was implemented after recognizing that some categories of fibrosis were spatially clustered across some sections. Such clustering resulted in oscillations in the estimate for the overall section from tile to tile, which would then dampen to a more accurate estimate after additional sampling.

Annotators were a team of six researchers involving a senior project scientist (BAJ), a postbaccalaureate researcher (NVE), and four undergraduate students (JMY, AG, AAN, JZC) trained initially by a board-certified neuropathologist (ESM). Additional training involved live interactive demonstrations, video recordings, illustrated manuals complete with examples of each fibrosis category, and live regular feedback sessions overseen by the project scientist.

### Deep-learning (DL) model for analysis of fibrosis

Training, validation, and test datasets were developed using the manual annotations. This included 29730, 6370, and 6372 images with fibrosis annotations for training, validation, and test dataset, respectively. A 6-layer U-Net like model was created with 4 output layers, one for each annotation category (papilla area, dense fibrosis, hyalinized fibrosis, and mineralized fibrosis). The DL model was trained using Keras (Chollet 2015) and TensorFlow v2.5 (Abadi et al 2016) on a NVIDIA RTX 3090 graphics processing unit.

Binary masks of labeled ChP papilla as well as masks of dense, hyalinized, and mineralized fibrotic regions were generated from the fibrosis-segmentation model output. Labeled papilla masks were refined using binary closing, opening, and fill operations to smooth out and remove holes from papilla segmentations followed by removal of objects smaller than 100 pixels in size (too small to be considered a papilla). Based on the masks, the amount of fibrosis as a percentage of papilla area was calculated on a per papilla basis.

Large lipid droplet coordinate data were obtained through a previously described DL model (Espericueta et al 2026) and mapped to the nearest ChP papilla based on a distance transform calculated from the previously generated papilla mask. Papilla-specific estimates of the percent of CPECs with large lipid droplets were then calculated and compared across fibrotic and non-fibrotic papilla using a paired-t test as implemented by the SciPy module in python (Virtanen et al 2020) or correlated against papilla-specific fibrosis percentages using Spearman correlations. All relevant code can be found in our GitHub repositories (https://github.com/mjneel/biondi & https://github.com/orgs/monukilab/repositories).

## RESULTS

### Classification of ChP fibrosis

After training to standardize definitions, a team of six researchers used customized software and polygon tracing tools to manually annotate randomly sampled portions of whole-slide images of H&E-stained sections taken from 81 specimens of postmortem human ChP (Fig. 1, center panels). The researchers separately traced areas of dense (little space between fibrils), hyalinized (glassy), and mineralized (purple concretions) fibrosis, also marking papilla from concavity to concavity. Their classifications were then used to train a DL model, which generally resulted in smoother and more reliable contours on the same images traced by the human annotators (Fig. 1, right panels).

### Agreement between manual and DL-model annotations

There was a significant correlation between the manual and DL-model estimates of both the percent of papillae affected across different cases (Fig. 2A) and the percent of papillary area affected by any level of fibrosis (Fig. 2B). Bland-Altman plots further established general agreement between the manual and DL-model estimates, although there was a tendency for DL model estimates to give lower values at lower amounts of fibrosis and there was a greater variance between the methods at higher amounts of fibrosis (Fig. 2C,D). When dense, hyalinized, and mineralized areas were independently evaluated, general agreements between the manual and DL-model analyses were also observed (Fig. S1).

**Fig. 2.**
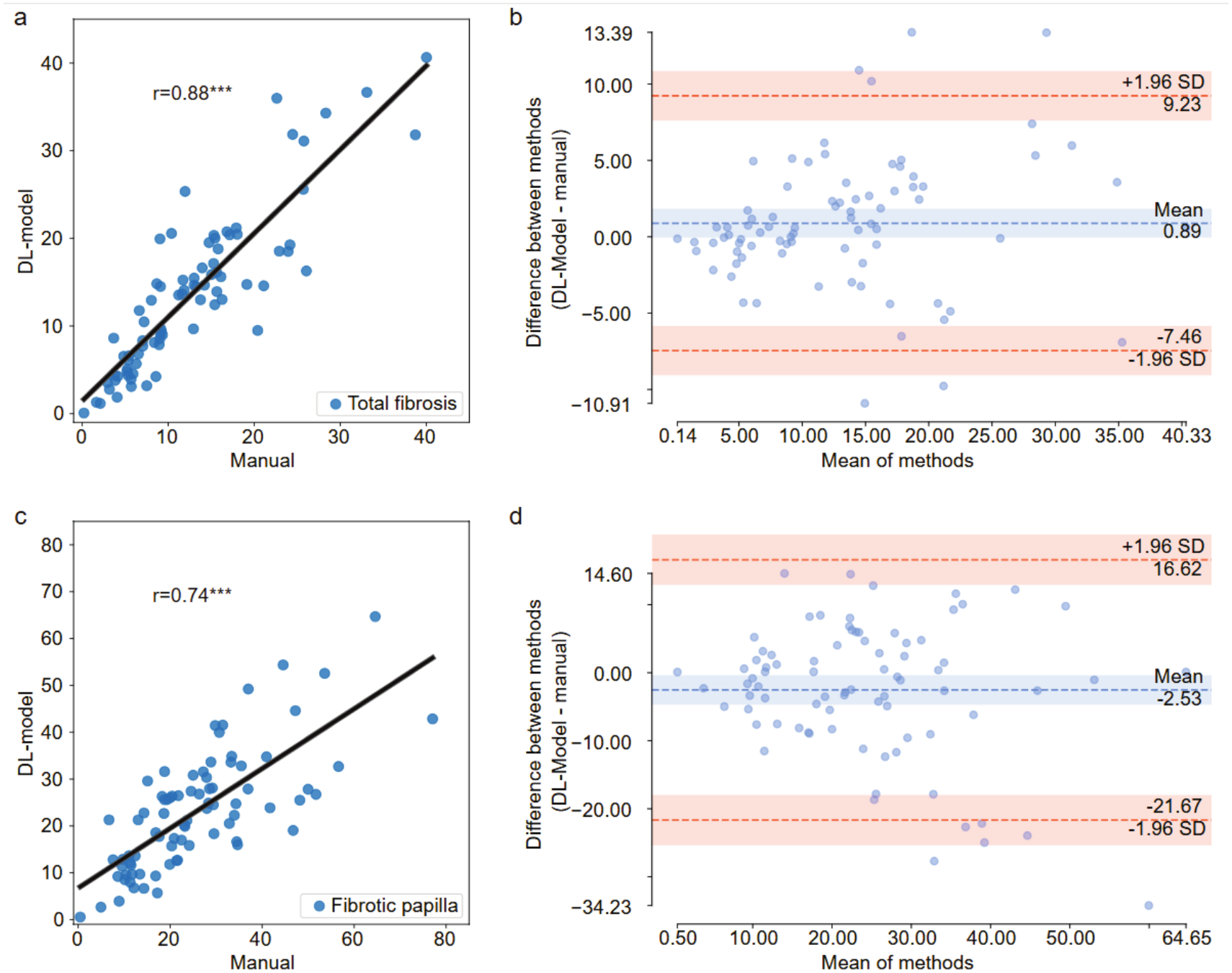
Fibrosis values determined by the DL model across entire whole slide images show good agreement with manual estimates. **a**) Correlation between manual and DL model values for the overall percent area of papillae affected by fibrosis. **b**) Bland-Altman plot of the agreement between manual and DL models in **a**. **c**) Correlation between manual and DL model values for the percent of papillae affected by any fibrosis. **d**) Bland-Altman plot of the agreement between manual and DL models in **c**.

### Increases in fibrosis with age

Given the good performance of the DL model, we evaluated overall fibrosis across entire whole slide images from the 199 cases in the UC Irvine repository for which we knew AD status. These included specimens taken at the UCI Medical Center as well as specimens from the UCI ADRC. The large scope of this analysis would not have been practical for manual annotators. Statistical analysis was conducted separately for cases across the lifespan and for cases over 65 years old, the latter approach to assess any possible association with Alzheimer’s disease. Both the percent of papillae affected by fibrosis and the percent of papillary area affected by fibrosis were significantly associated with age (Table 1). No association was found for sex or AD, but there was a significant interaction between age and the source of tissue for the percent of papilla area affected (Table 1). When plotted as a function of age, the percent affected area for each of the two sources appeared to increase linearly, but with different slopes (Fig. 3A). A plot of age versus the percent of papilla affected by any fibrosis could be fit adequately with a single line regardless of the tissue source (Fig. 3B).

**Fig. 3.**
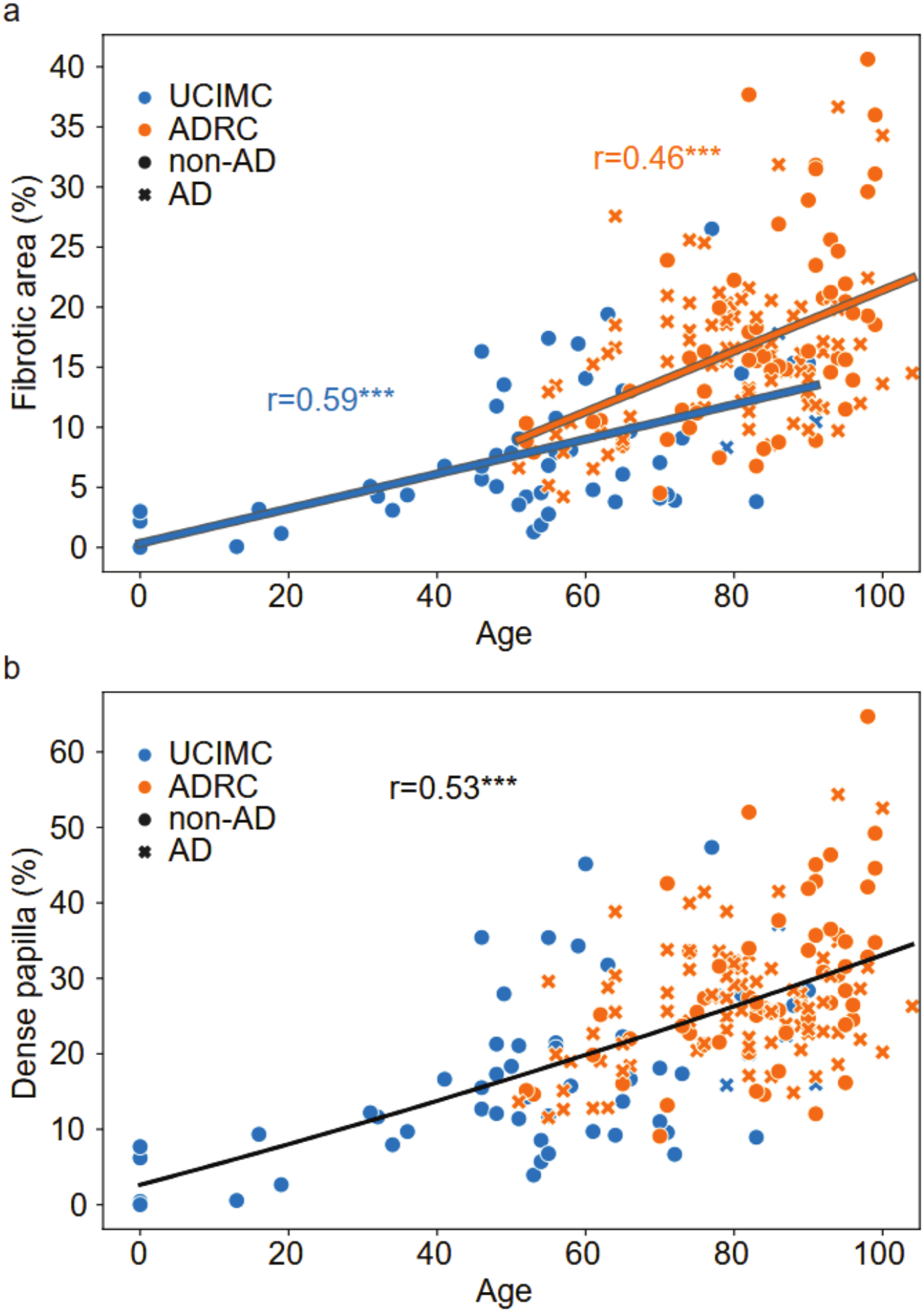
Fibrosis increases with age. **a**) DL model characterization of the percent papillary area affected by fibrosis as a function of age. Data points were fit to two lines representing the two different tissue sources, which showed a significant interaction with age in statistical analyses (Table 1). **b**) DL model characterization of the percent of papillae affected by fibrosis as a function of age.

**Table 1.** P values from ANCOVA analyses of overall fibrosis.

| variable | All cases (n=199) |  | Cases ≥ 65 years (n=141) |  |
|---|---|---|---|---|
|  | % Papilla | % Area | % Papilla | % Area |
| AD |  |  | 0.29 | 0.51 |
| Sex | 0.31 | 0.28 | 0.36 | 0.38 |
| Age | < 0.001*** | < 0.001*** | < 0.001*** | < 0.001*** |
| Source | 0.54 | 0.17 | 0.24 | 0.42 |
| Age * Source | 0.21 | 0.013* | 0.42 | 0.72 |

### Spatial clustering of fibrotic papillae

DL model-based annotation of fibrosis allowed us to investigate the possible spatial clustering of affected papillae across whole slide images. Each papilla was evaluated for the percentage of papillae within 700 microns that contained any fibrosis. Three examples of the distribution are illustrated in Fig. 4. Discrete areas of high and low fibrosis were observed, which was particularly obvious for case #499605. Statistical analysis employed Kolmogorov-Smirnov tests, in which histograms of local density for each case were compared to 1000 Monte Carlo simulations of random distributions. Statistically significant (p < 0.05) spatial clustering was found for 195 out of 204 cases. The degree of spatial clustering, as reflected in the Kolmogorov-Smirnov D-statistic, was significantly and positively correlated with age (Fig. S2).

**Fig. 4.**
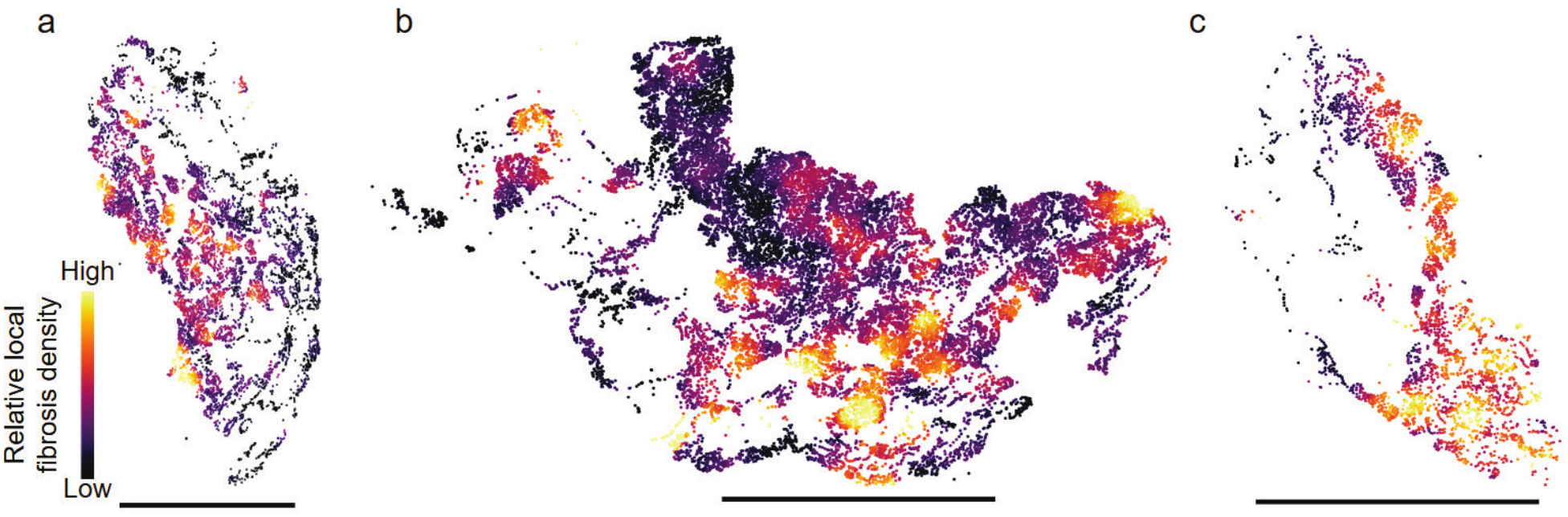
Fibrosis is spatially clustered within the ChP. Scatter plots show the location of annotated ChP papillae in a single section of each of three cases (**a-c**; cases 363485, 499605, & 335909). Each point is color coded to represent the average percent area affected by fibrosis in papillae within 700 μm. Scale bars denote 1 cm.

### CPECs associated with fibrotic papillae have fewer large lipid droplets

The same H&E-stained sections used for analysis of fibrosis also had been used for analysis of CPECs containing large lipid droplets (Espericueta et al 2026), which enabled a detailed analysis of the spatial relationship between the two age-related changes to the ChP. Large lipid droplet coordinate data were mapped to the nearest ChP papilla, and papilla-specific values were then calculated and compared across fibrotic and non-fibrotic papillae. CPECs with large lipid droplets were significantly less abundant in fibrotic papillae (p < 0.001, paired t-test, n=192) (Fig. 5).

**Fig. 5.**
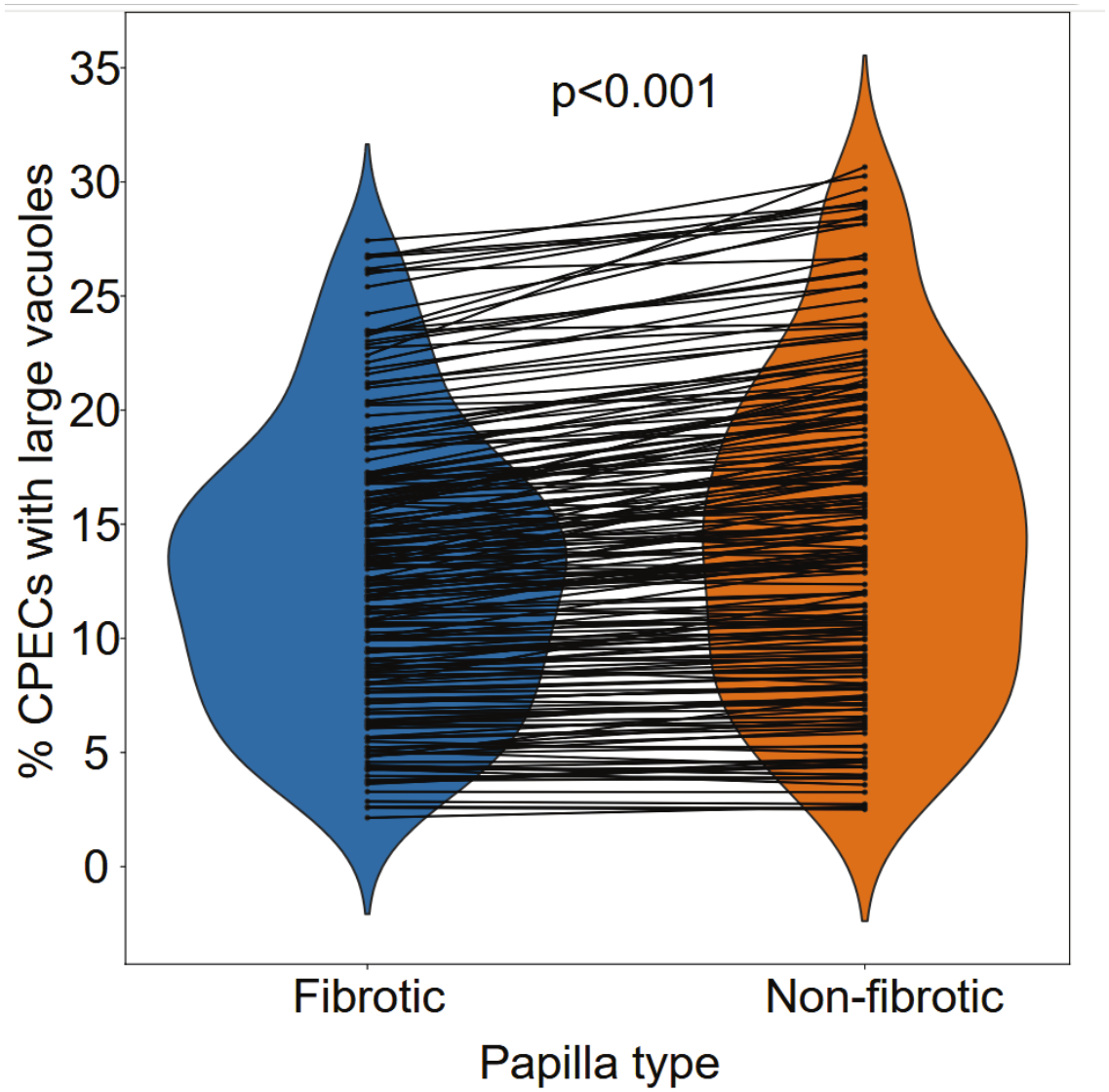
CPECs surrounding fibrotic papillae have fewer large lipid droplets. Violin plots comparing the percentage of CPECs with large lipid droplets between fibrotic and non-fibrotic papilla. Lines between the violin plots connect data from the same case. A paired t-test showed the differences to be significant (p< 0.001).

### All stages of fibrosis increase with age

In addition to overall fibrosis, we separately analyzed areas of fibrosis that we classified into non-overlapping, increasingly affected categories: dense, hyalinized, and mineralized (Fig.1, Fig S1). The area affected by each of the categories was found to be significantly associated with age (Fig. 6, Table 2). In addition, hyalinized and mineralized levels showed a significant interaction with the source of the tissue.

**Fig. 6.**
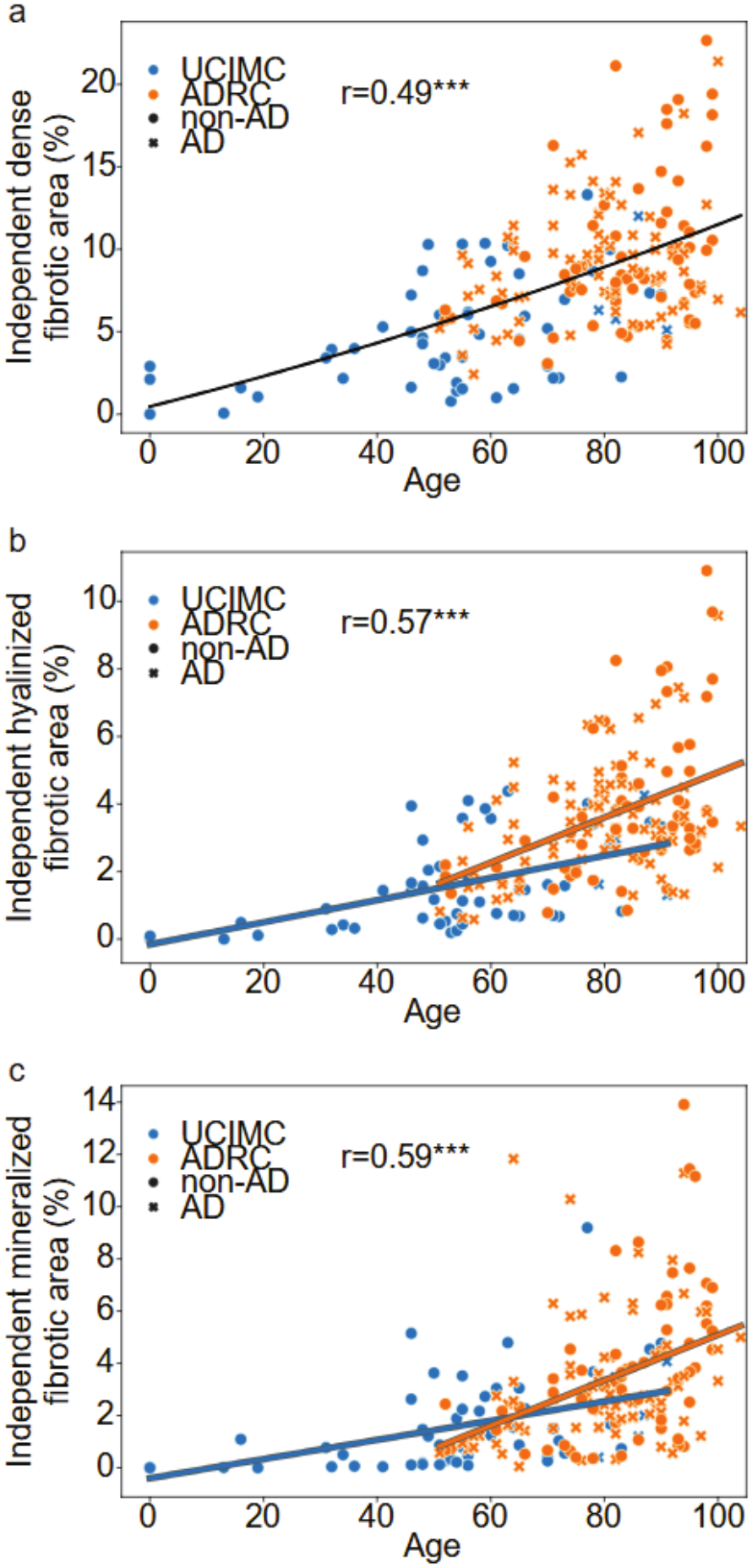
All stages of fibrosis increase with age. Fibrotic areas annotated as being only **a**) dense (but neither hyalinized nor mineralized), **b**) hyalinized (but not mineralized), or **c**) mineralized were all found to increase with age. Hyalinized and mineralized fibrosis were fit with two lines representing the two different tissue sources, which showed a significant interaction with age in statistical analyses.

**Table 2.** P values from ANCOVA analyses of stages of fibrosis.

| Variable | All cases (n=199) |  |  | Cases ≥ 65 years (n=141) |  |  |
|---|---|---|---|---|---|---|
|  | Dense | Hyalinized | Mineralized | Dense | Hyalinized | Mineralized |
| AD |  |  |  | 0.79 | 0.80 | 0.24 |
| Sex | 0.035* | 0.39 | 0.32 | 0.073 | 0.40 | 0.36 |
| Age | <0.001*** | <0.001*** | <0.001*** | 0.004** | <0.001*** | <0.001*** |
| Source | 0.86 | 0.067 | 0.01* | 0.52 | 0.20 | 0.86 |
| Age*source | 0.30 | 0.005** | 0.004** | 0.82 | 0.42 | 0.98 |

Specimens from the ADRC, which involved older cases overall, displayed a steeper age dependency than specimens from the UCI Medical Center autopsy service. There was a significant sex effect for the dense category, with males being more affected than females. No association with Alzheimer’s disease was detected for cases aged 65 years or older.

### All stages of fibrosis are associated with decreased large lipid droplets in associated CPECs

Across every papilla within each case, we correlated the percentage of associated CPECs containing large lipid droplets with the area of the papilla that was affected by each category of fibrosis. Fig. 7 shows the Pearson correlation coefficients obtained for each case and fibrosis category. The great majority of cases (88%, 89%, and 88% of dense, hyalinized, and mineralized fibrosis datapoints, respectively) resulted in negative correlations, and one-sample t-tests found that the correlation coefficients across cases were significantly different than zero (P<0.001 for each category).

**Fig. 7.**
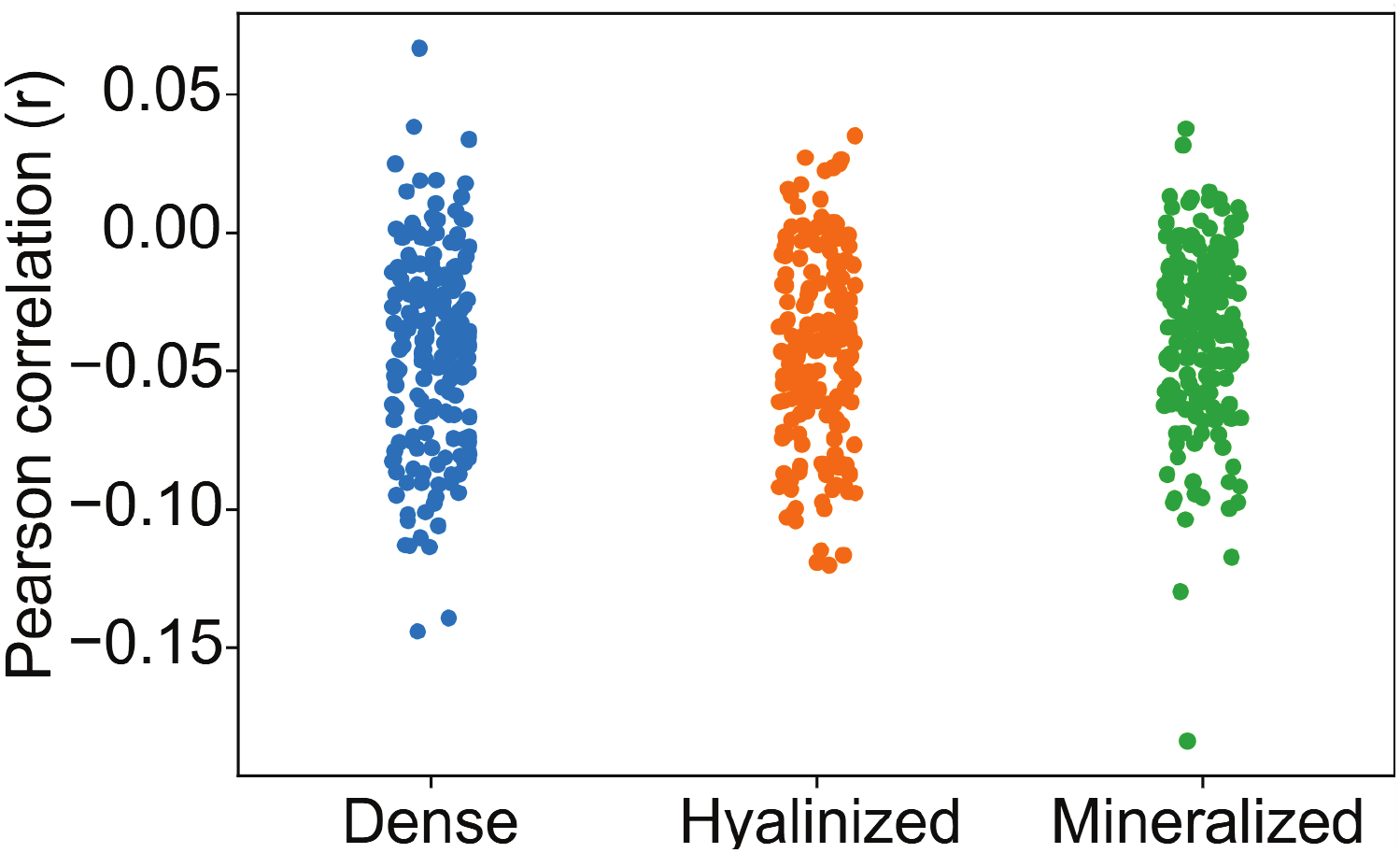
The percentage of CPECs with large lipid droplets was negatively correlated with the area of fibrosis in their associated papillae at all stages of severity. Correlations were mostly negative, and one-sample t-tests showed significant differences from zero for all three stages.

### Individuals with more nodular stromal fibrosis had fewer CPECs with large lipid droplets

Given the papilla-by-papilla negative association between stages of fibrosis and large lipid droplets within a case, we also analyzed associations across cases to determine if individuals with more fibrosis displayed fewer CPECs with large lipid droplets. As shown in Table 3, a significant negative association was indeed detected for the papilla area occupied by any fibrosis when corrected for age, sex, and tissue source. Significant associations of large lipid droplets with age and tissue source were also detected in this analysis. Statistical analysis of independently defined stages of fibrosis did not reveal further significant associations with large lipid droplets.

**Table 3.**
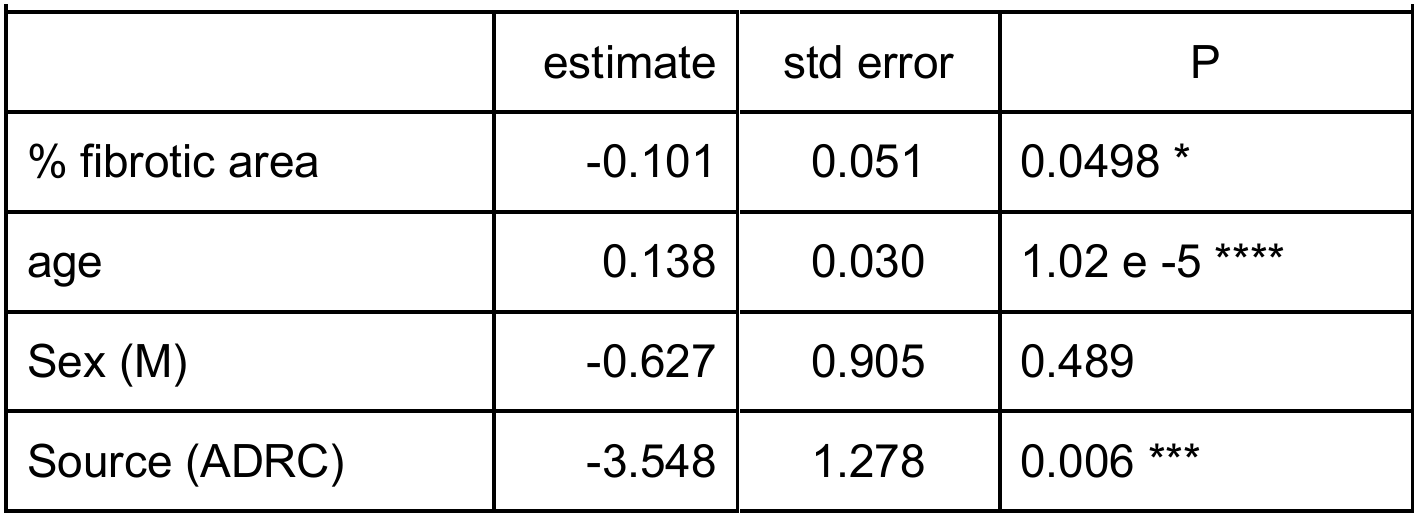
Large lipid droplets across donors (ANCOVA, n=203)

## DISCUSSION

We have replicated prior findings of age-associated increases in the percentages of papillae affected by fibrosis in the postmortem human ChP (Dunn & Kernohan 1955, Prineas et al 2016), refining the work by assessing areas affected as well as by classifying independent stages of fibrosis that reflect increasing severity. We found that nodular stromal fibrosis is spatially clustered in the choroid plexus and that affected papillae are spatially associated with epithelial cells containing fewer large lipid droplets. Additionally, individuals whose choroid plexus is more affected by nodular fibrosis tended to have fewer large lipid droplets.

Despite the negative association between fibrosis and large lipid droplets, both fibrosis and large lipid droplets increase with age (Espericueta et al 2026), which may seem paradoxical. However, the effect of fibrosis on large lipid droplets appears to be local, indicating that the positive effect of age on large lipid droplets is larger overall than is the negative local fibrosis effect.

### Increased ChP fibrosis may be related to changes detected using in vivo imaging

MRI has revealed age-dependent decreases in ChP water mobility and capillary permeability that may be related to stromal fibrosis (Bouzerar et al 2013, Alisch et al 2021). Computed tomography (CT) of human ChP has uncovered aging-associated increases in calcification (Macpherson & Matheson 1979, Bersani et al 1999, Yalcin et al 2016), which may be related to the age-related increases in mineralized papillary fibrosis we have observed, although psammoma bodies in the ChP stroma (Alcolado et al 1986, Jovanovic et al 2007) also contribute to the CT signal.

### Spatial clustering of fibrosis in the human ChP

We have found that fibrotic papillae occur in spatial clusters across the ChP. These findings suggest that different parts of the ChP are affected by age differently and/or at different rates. Fibrosis is associated with markers of ChP inflammation (Prineas et al 2016), which may point to localized responses to spontaneous injury, infection, or mutation. Our finding of a negative association between fibrotic papillae and CPECs with large lipid droplets, which also increase with age (Espericueta et al 2026), is consistent with a distinct spatial distribution of these two age-related changes.

### Negative association between fibrosis and large lipid droplets in CPECs

It seems likely that fibrosis impairs transport of materials from blood to CPECs, including the water that makes up most of the CSF and the immune cells transiting the blood-CSF barrier, possibly contributing to age-related decreases in CSF production (May et al 1990, Preston 2001, Nizari et al 2025).

We now have shown that fibrotic papillae are associated with relatively fewer CPECs containing large lipid droplets compared to non-fibroic papillae. The prevalence of CPECs with large lipid droplets in postmortem material is significantly correlated with body mass index at time of death, and CPECs derived from human stem cells exhibit fatty acid uptake into large droplets, especially when oleate is added to the culture medium (Espericueta et al 2026).

We have proposed that fatty acids carried by albumin are taken up and stored by CPECs (Espericueta et al 2026) for the CPECs’ own considerable energetic needs as well as for packaging and release as lipoproteins into CSF to support the brain’s need for lipid (Qadir et al 2026). This process likely involves the secretion of lipase from the basal aspect of CPECs, which would then diffuse through stroma to release fatty acid from lipoproteins at the capillaries, followed by diffusion of albumin-bound fatty acid back through stroma to reach the CPECs (Song et al 2025). Nodular stromal fibrosis likely produces a barrier to this process in both directions such that fatty acids do not reach CPECs in sufficient concentration to result in large lipid droplets, thus potentially interfering with a pathway for lipid transport to the brain as well as depriving CPECs of an energy source that supports CSF production and conditioning. Alternatively (or additionally), local fibrosis may continuously stress associated CPECs metabolically, causing them to utilize and deplete their lipid stores.

## ACKNOWLEDGEMENTS

We thank UCI neuropathologists and technicians at the Experimental Tissue Resource and UCI ADRC Neuropathology Core for their assistance in procuring and processing tissue, staining, and slide scanning. This work was supported by the following NIH awards: P50AG016573 and P30AG066519 (UCI ADRC), T32AG073088 (N.V.E. and M.J.N.), T32AG000096 (M.J.N.), and R21MH109036 and R21AG064640 (E.S.M.). We are greatly indebted to the tissue donors and their families for making this research possible.

---

**Fig. S1.**
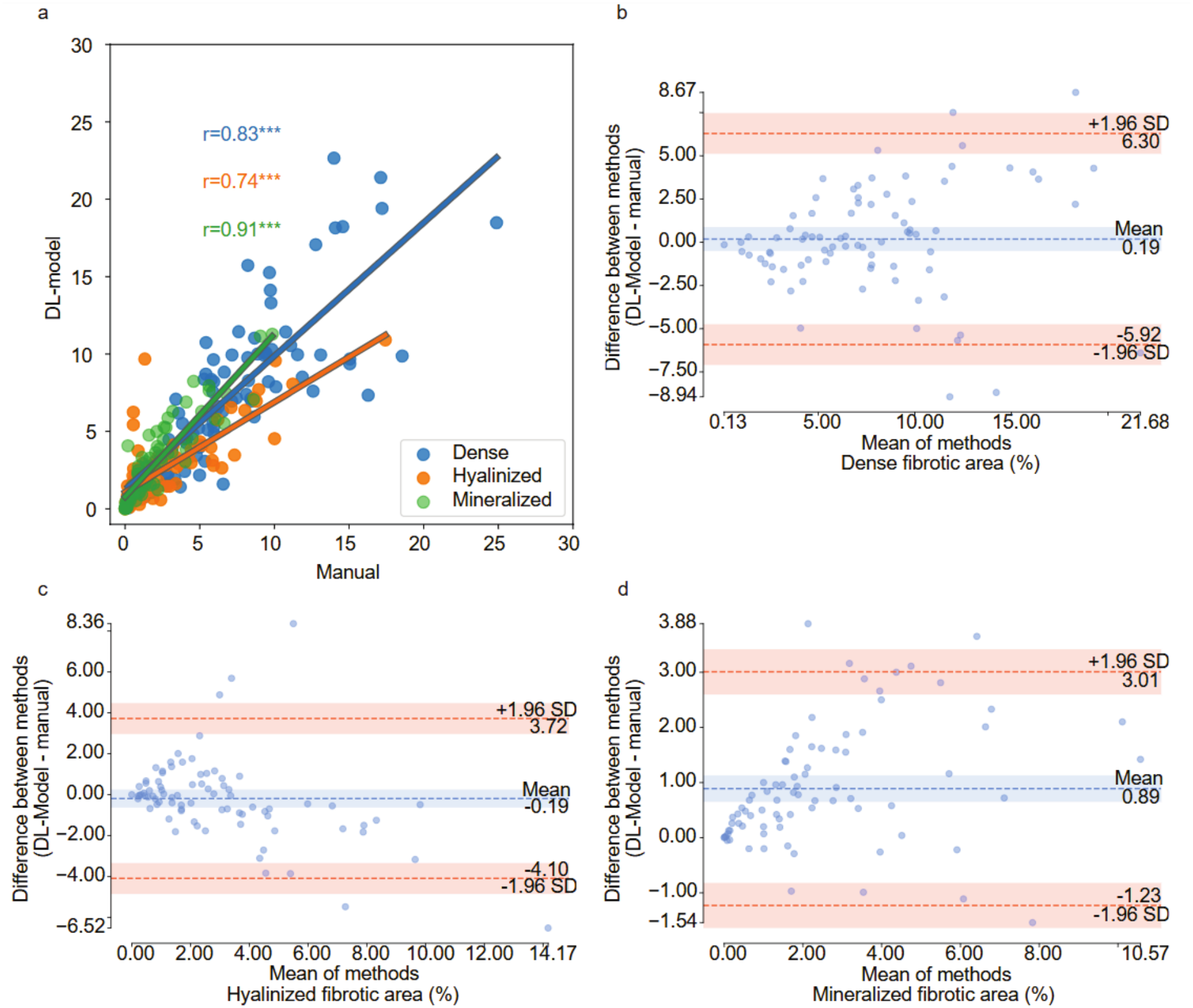
The DL-model recapitulates the results of manual annotations at each stage of fibrosis. **a**) Correlations between manual and DL model values for the percent area affected by each level of fibrosis. **b-d**) Bland-Altman plots of the agreement between manual and DL model results shown in **a**:.

**Fig. S2.**
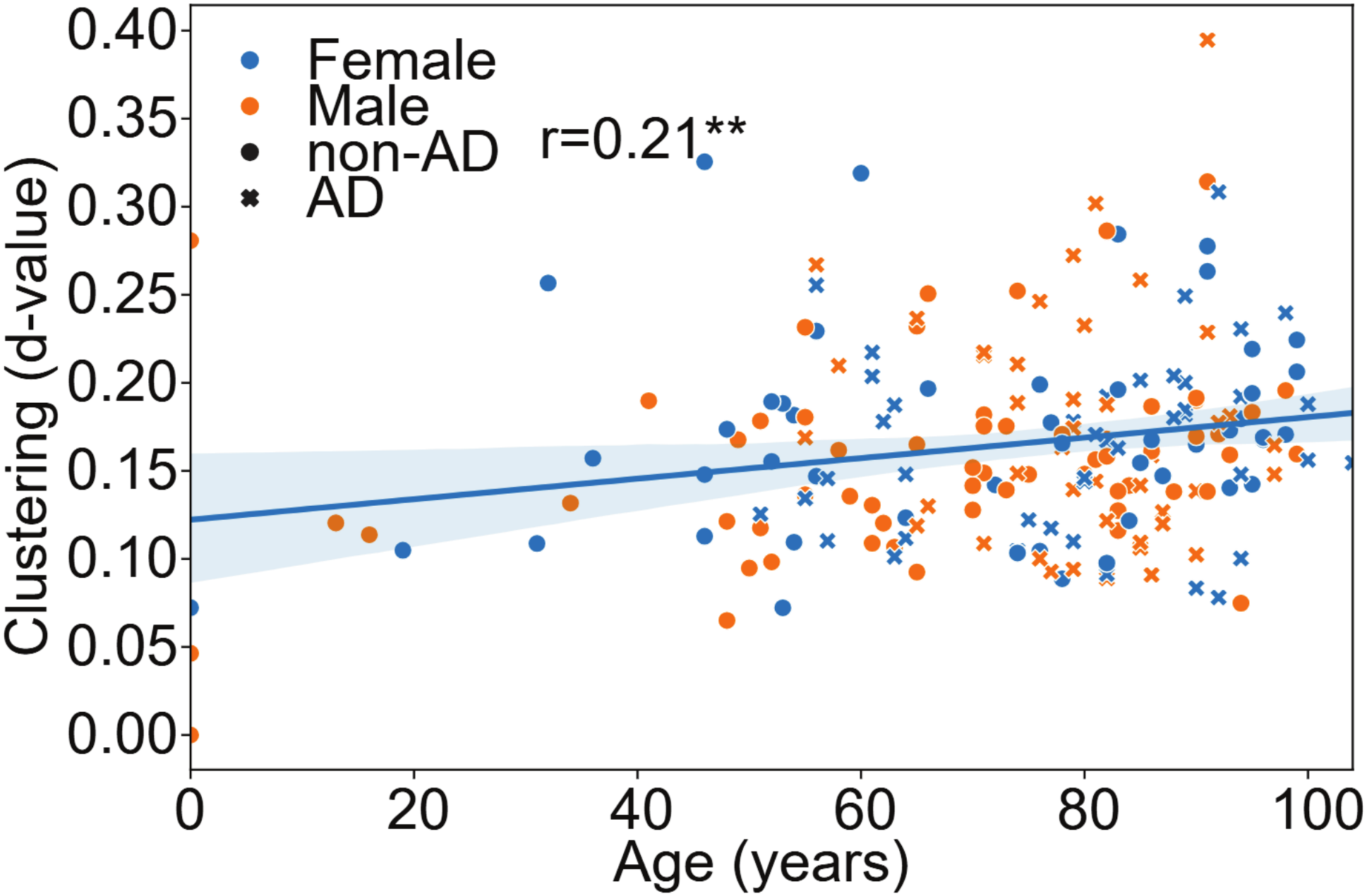
Fibrosis clustering correlates with age. The scatterplot shows DL-generated d-values from Kolmogorov–Smirnov statistical tests. The trendline is the result of a linear regression. Blue-shaded areas indicate 95% confidence intervals. The R-value is a Spearman correlation coefficient.

